# Glucagon Receptor Signaling at White Adipose Tissue Does Not Regulate Lipolysis

**DOI:** 10.1101/2022.03.20.485051

**Authors:** Anastasiia Vasileva, Tyler Marx, Jacqueline L. Beaudry, Jennifer H. Stern

## Abstract

**Objective:** Although the physiologic role of glucagon receptor signaling in the liver is well defined, the impact of glucagon receptor (Gcgr) signaling at white adipose tissue (WAT) continues to be debated. While numerous studies propose glucagon stimulates WAT lipolysis, we lack evidence that physiological concentrations of glucagon regulate WAT lipolysis. Glucagon receptor antagonists are proposed as a treatment to lower blood glucose in people with type 2 diabetes, yet concerns on how these treatments may affect lipid homeostasis have led to questions regarding the potential safety and efficacy of such therapeutics. Tight regulation of adipose tissue lipolysis is critical for whole body lipid homeostasis. In turn, we used WAT *Gcgr* knockout mice to determine if glucagon regulates lipolysis at WAT in the mouse.

**Methods:** We assessed the effects of fasting and acute exogenous glucagon administration in wildtype C57BL/6J and *Gcgr* ^Adipocyte+/+^ vs *Gcgr*^Adipocyte-/-^ mice. Using an *ex vivo* lipolysis protocol, we further examined the direct effects of glucagon on physiologically (fasted) and pharmacologically stimulated lipolysis.

**Results:** Adipocyte Gcgr expression did not affect fasting induced lipolysis or hepatic lipid accumulation in lean or diet induced obese (DIO) mice. Acute glucagon administration did not affect serum non-esterified fatty acids (NEFA), leptin, or adiponectin concentration, but did increase serum glucose and FGF21, regardless of genotype. Glucagon did not affect *ex vivo* lipolysis in explants from either *Gcgr* ^Adipocyte+/+^ or *Gcgr*^Adipocyte-/-^ mice. *Gcgr* expression did not affect fasting-induced or isoproterenol-stimulated lipolysis from WAT explants. Moreover, glucagon receptor signaling at WAT does not affect body weight or glucose homeostasis in lean or DIO mice.

**Conclusions:** We have established that glucagon does not regulate WAT lipolysis, either directly or indirectly. Unlike the crucial role of hepatic glucagon receptor signaling in maintaining glucose and lipid homeostasis, we observed no metabolic consequence of WAT glucagon receptor deletion.

## 1. Introduction

Glucagon plays a critical role in the maintenance of glucose (1) and lipid (2) homeostasis during fasting, primarily by stimulating glycogenolysis, gluconeogenesis, fatty acid oxidation and inhibiting *de novo* lipogenesis at the liver. Although the liver is the main site of glucagon action and receptor expression, glucagon receptors have been detected to a far lesser extent in mouse (3), rat (4), and human (5, 6) white adipose tissue (WAT). Whether glucagon receptor signaling at WAT plays a role in metabolic homeostasis is not clear. Glucagon has been shown to directly activate hormone-sensitive lipase in rat epididymal fat pads (7) and stimulate lipolysis in isolated rat (8-12) and human (6) white adipocytes. Yet, clinical studies demonstrate that physiological levels of glucagon have no effect on adipose tissue lipolysis in people with (13) or without (13, 14) diabetes.

Obese, insulin resistant humans (15-17) and mice (17) hyper-secrete glucagon in the fed state, exacerbating hyperglycemia and encouraging the development of type 2 diabetes (T2DM). Suppression of glucagon signaling improves glycemic control and decreases hyperglycemia in people with insulin resistance (18, 19). Accordingly, the development of glucagon receptor antagonists and antibodies are currently being pursued as a glucose-lowering therapeutic for patients with T2DM (20-22). Suppression of glucagon signaling is similarly effective at improving glycemic control in hyperinsulinemic/ insulin resistant mice (23, 24). Accordingly, the mouse is an informative model to assess the potential beneficial responses to glucagon receptor manipulation. Despite improving the regulation of blood glucose, reported increases in hepatic triglyceride content, plasma total cholesterol (25), and liver enzymes (25, 26) have led to questions regarding the safety and efficacy of such therapeutics. Global glucagon receptor knockout increases fasting serum non-esterified fatty acid (NEFA) concentration, triglyceride (TAG) concentration, and hepatic TAG secretion in mice (2). Thus, the mouse is also an ideal pre-clinical model to assess the potential negative consequences of glucagon receptor antagonism.

The deleterious effects of glucagon receptor antagonism on lipid homeostasis may be due to a lack of glucagon receptor signaling at the hepatocyte or the adipocyte. Tight regulation of adipose tissue lipolysis is a critical factor in the maintenance of whole body lipid homeostasis. Suppression of WAT lipolysis decreases hepatic lipid accumulation (27), while elimination of insulin-mediated suppression of WAT lipolysis leads to increased hepatic lipid accumulation (28, 29), predisposing to hepatic insulin resistance and the development of liver disease. We set out to understand if glucagon signaling at WAT is involved in the regulation of WAT lipolysis and potential alterations in hepatic lipid concentration. Given that efforts to develop pharmacological inhibitors of glucagon receptor signaling as a therapy for hyperglycemia in T2DM continue (20, 21, 30), it is critical to understand the potential consequences of blocking glucagon receptor signaling at the adipocyte.

## 2. Materials and Methods

### 2.1 Mice

All mice were maintained on a 12-hour light/12-hour dark cycle and housed with 3-5 mice per cage until 1 week prior to study initiations, at which time animals were individually housed. Mice were housed with sani-chip bedding (7090 Teklad). All studies were approved by The University of Texas Southwestern and The University of Arizona Institutional Animal Care and Use Committees. All mice were provided ad libitum access to food (2016 Teklad Global 16% Protein Rodent Diet, 12% kcal from fat; Envigo, Indianapolis, IN) and water unless fasting or alternative diets are specified.

Terminal glucagon responsivity studies were performed in male wildtype C57BL/6J mice that were bred in-house at UT Southwestern Medical Center. *Ex vivo* lipolysis assays performed in wildtype male C57BL/6J mice were purchased from Jackson laboratories.

Adiponectin-rtTA (Apn-rtTA) (31) and floxed *Gcgr* (1) mice were generated as previously described. The TRE-Cre mouse was purchased from Jackson Laboratories (Strain # 006234; Jackson Laboratories, Bar Harbor, ME, USA) Gcgr^Adipocyte-/-^ (adipocyte-specific Gcgr knockout; Apn-rtTA^+/-^, TRE-Cre^+/-^, Gcgr^F/F^) mice and littermate Gcgr ^Adipocyte+/+^ controls (Apn-rtTA^+/-^, TRE-Cre^-/-^, Gcgr^F/F^) were generated by crossing male Apn-rtTA-/-, TRE-Cre+/-, Gcgr^F/F^ mice to female Apn-rtTA+/-, TRE-Cre-/-, Gcgr^F/F^ mice. Mice were fed a standard chow diet (2016 Teklad Global) until approximately 8 weeks of age, at which point a chow (S4107, Bio-Serv, Flemington, NJ) or high fat (60% energy from fat; Bio-Serv S5867) diet containing 600mg/kg doxycycline (DOX) was provided to initiate adipocyte-specific deletion of the glucagon receptor gene. Chow fed mice were maintained on DOX for 4 weeks prior to the initiation of studies. High fat diet fed mice were maintained on high fat DOX diet for 14 weeks to induce obesity.

### 2.2 Terminal Glucagon Responsivity Tests

To initially examine the effect of exogenous glucagon administration on adipose tissue lipolysis, male wildtype C57BL/6J mice were fasted for either 4 or 16 hours prior to IP injection with either saline or glucagon (5μg/kg BW; Eli Lily and company, Indianapolis). Mice were sacrificed at either 0 (saline group only), 15, 30, or 60 minutes after injection by decapitation after bell jar exposure to isoflurane anesthesia. Trunk blood was immediately collected and allowed to clot prior to centrifugation at 3,000 x g for 30 minutes. Serum was collected, aliquoted, and immediately frozen at -80°C. Tissues were collected immediately, rinsed with phosphate-buffered saline and flash frozen in liquid nitrogen and stored at -80C until analysis. Terminal glucagon responsivity tests in Gcgr ^Adipocyte+/+^ and Gcgr^Adipocyte-/-^ mice were performed identically, with mice sacrificed 15 minute after injection.

### 2.3 Crossover glucagon responsivity tests

Male Gcgr ^Adipocyte+/+^ and Gcgr^Adipocyte-/-^ mice were fasted for either 4 or 16 hours prior to IP injection with either saline or glucagon (5μg/kg BW; Eli Lily and company, Indianapolis). 30 minutes after injection, tail blood was collected using a capillary tube and allowed to clot prior to centrifugation at 3,000 x g for 30 minutes. Serum was collected, aliquoted, and immediately frozen at -80°C. Mice were allowed to recover for 3 days before the study was repeated in a crossover fashion.

### 2.4 *Ex vivo* lipolysis

*Ex vivo* lipolysis studies were performed in male 16-18 week old mice. Mice were fasted for 4 or 16 hours prior to being sacrificed by decapitation after bell jar exposure to isoflurane anesthesia. *Ex vivo* lipolysis was assayed as previously described (32, 33). Briefly, gonadal adipose tissue was collected immediately after euthanasia and washed with phosphate-buffered saline prior to mincing tissue into ∼20mg pieces. Lipolysis was assessed in triplicate for each treatment within a mouse. Explants were then incubated for 1 hour in Krebs-Ringer buffer (12 mM HEPES, 121 mM NaCl, 4.9 mM KCl, 1.2 mM MgSO4, and 0.33 mM CaCl2, 3 mM glucose containing 2% fatty acid free bovine serum albumin) at 37°C, then transferred to Krebs-Ringer buffer containing the following treatments: 10µM isoproterenol, 10µM forskolin, insulin (20nM, 200nM, and 200nM), glucagon (0.1nM, 1nM, 10nM, and 100nM), or control (Krebs-Ringer only). After 1h incubation, explants were removed and stored at -80°C until analysis for total protein content. Explants were sonicated in 0.1 M Phosphate Buffered Saline, pH 7.4 (PBS), then centrifuged for 10 minutes at 13,000 x g at 4°C. The top lipid layer was carefully removed prior to transferring supernatant to a fresh tube. Total protein in the supernatant was assayed using a colorimetric assay (Pierce™ BCA Protein Assay Kit, # 23225). Media was collected and stored at -80°C until analysis for NEFA content via colorimetric assay (999-34691, 995-34791, 991-34891, and 993-35191, Wako Diagnostics USA). Leptin content in media was analyzed by ELISA (Cat. # EZML-82K, Millipore Sigma, Danvers, MA).

### 2.5 Oral glucose tolerance testing

After a 4h fast, we gave individually housed mice an oral gavage of D-glucose (2.5g/kg; Fisher) and assessed blood glucose by glucometer (9556c, Bayer, Leverkusen, Germany). Blood was collected by tail nick at 0, 15, 30, 60, 90, and 120 minutes following glucose gavage. Blood for serum insulin (glucose stimulated insulin secretion) was collected from the tail vein prior to and 15 minutes following glucose administration.

### 2.6 Oral lipid clearance testing

After an overnight 16h fast, olive oil (10µl/g body weight) (34) was gavaged into individually housed mice. Oral lipid clearance testing began at 9am. Blood for serum triglyceride was collected via tail vein at 0, 30min, 1, 2, 4, 6, 8, and 10 hours following olive oil gavage.

### 2.7 Tyloxapol stimulated triglyceride

After a 4h fast, individually housed mice were injected with 300 mg/kg Triton WR-1339 (Tyloxapol; Sigma-Aldrich) in 0.9% saline via the tail vein (35) to inhibit lipoprotein lipase activity. Studies began at 1pm and blood for serum triglyceride was collected via tail vein at 0, 30, 60, 90, and 120 minutes following injection.

### 2.8 Serum analyses

Commercially available enzyme-linked immunosorbent assays were used to assess hormones in serum and media (Glucagon : Cat. # 10-1271-01, Mercodia, Uppsala, Sweden; Insulin: Cat. # 80-INSMSU-E10, Alpco, Salem, NH; FGF21: Cat. # EZRMFGF21-26K, Millipore Sigma, Danvers, MA; Leptin: Cat. # EZML-82K, Millipore Sigma, Danvers, MA;and Adiponectin: Cat. # EZMADP-60K, Millipore Sigma, Danvers, MA). Serum glucose, NEFA, TAG, and total cholesterol concentrations were analyzed by an enzymatic colorimetric assay (Glucose: Cat. # G7519, Pointe Scientific Inc., Canton MI; NEFA: Cat. # 999-34691, 995-34791, 991-34891, and 993-35191, Wako Diagnostics USA; TAG: Cat. # T7531, Pointe Scientific Inc., Canton, MI; and total cholesterol: Cat # TR13421, Thermo Scientific, Middletown, VA).

### 2.9 Tissue Analyses

#### 2.9.1 RNA isolation and gene expression

RNA was isolated using TRIzol™ Reagent (Thermo Fisher Scientific, Waltham, MA). Phenol was eliminated using the water saturated butanol and ether method of Krebs, Fischaleck, & Blum (36). Reverse transcription was performed using Verso cDNA synthesis kit (Thermo Scientific, Inc., Waltham, MA), and RT-qPCR performed using PowerUp™ SYBR™ Green Master Mix on the Applied Biosystems QuantStudio 6 Flex Real-Time PCR System (Applied Biosystems™, Foster City, CA). LinReg PCR analysis software was used to determine the efficiency of amplification from raw CT data (37). ACTβ served as the reference gene for calculating fold change in gene expression using the efficiency^ΔΔCt^ method (38). Gcgr mRNA was detected using Taqman probes Mm00433546_m1 and m00433550_g1 (ThermoFisher, Waltham, MA). Mouse primer sequences for all other genes for real-time PCR are presented in Table 1.

**Table 1.**
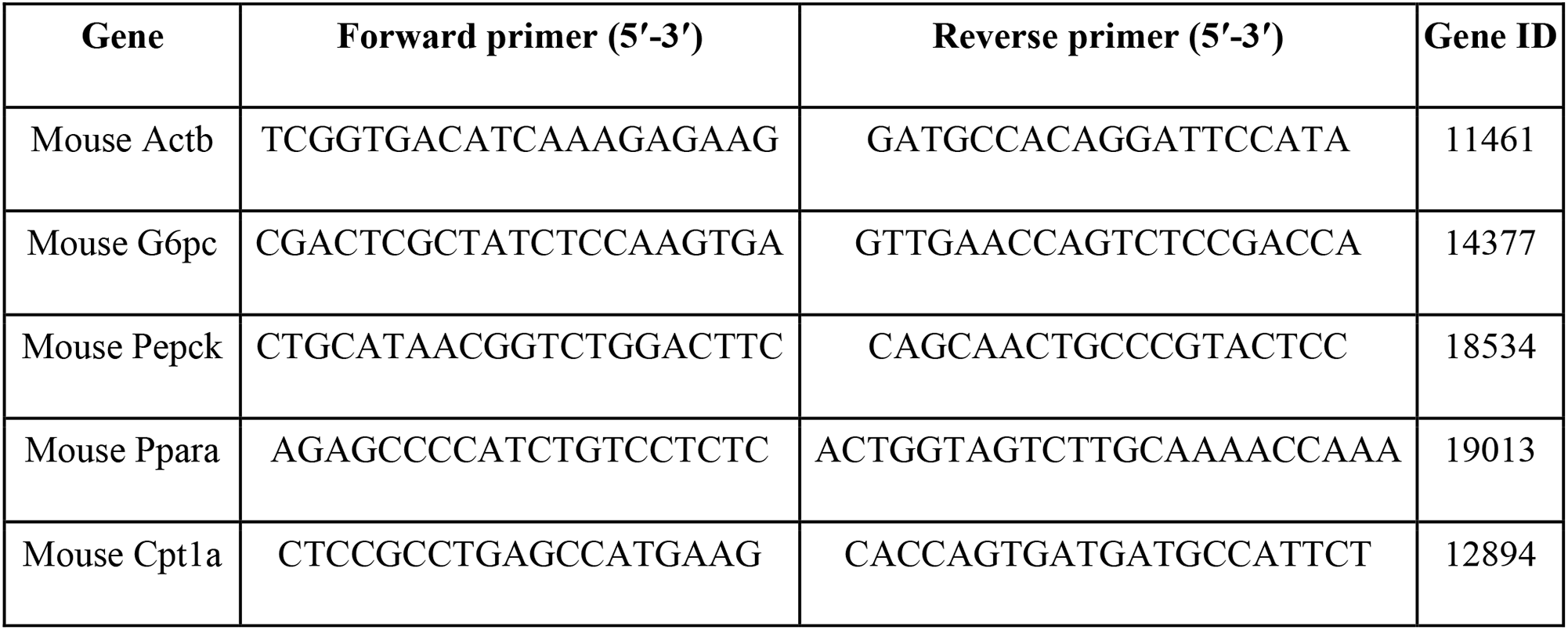
List of primer sequences for RT-PCR

#### 2.9.2 Hepatic lipid content

Livers were powdered with a liquid nitrogen cooled mortar and pestle to ensure a homogenous sample. Briefly, 10-20 mg of powdered liver samples were weighed and sonicated in 100µL PBS. 1 mL of 100% ethanol was added to each sample and vortexed for 20 minutes then centrifuged at 16,000xg at 4°C (39). Supernatant was then transferred to a fresh tube for analysis of liver triglycerides (Cat. # T7531, Pointe Scientific Inc., Canton, MI). Total hepatic triglyceride content was calculated as mg/g tissue.

### 2.10 Immunohistochemistry and imaging of pancreata

Immediately after mice were sacrificed, whole pancreata were collected into 4% paraformaldehyde and fixed for 24 hours. Tissues were then transferred to a 50% ethanol solution prior to paraffin embedding. Paraffin embedding and tissue sectioning was performed by the Molecular Pathology Core Facility at UTSW. Slides were then deparaffinized with xylene (3minute incubation) and rehydrated using graded concentrations of water:ethanol (3-minute incubations at 0:100, 5:95, 30:70, and 50:50), followed by a 3 minute incubation in PBS. Slides were boiled in antigen retrieval solution (10mM Sodium Citrate with 0.05% Tween-20, pH 6.0) for 12 minutes, then allowed to cool at room temperature for 30 minutes. Subsequently, immunohistochemistry was performed. Briefly, slides were washed 3 × 1 minute in PBS before exposing to a blocking solution containing TBST (TBS with 0.1% Tween-20) plus 20% AquaBlock (Cat. # ab166952, Abcam USA) for 30 minutes. Following blocking, slides were incubated overnight at 4°C in primary antibodies (1:500 dilution in blocking solution) for glucagon (Abcam, cat. # ab10988) and insulin (Cat. # A0564, Dako, Carpinteria, CA). Each slide contained a negative control section incubated only in blocking solution. Slides were next washed 3 times for 5 minutes in PBST (PBS with 0.05% Tween-20). Slides were then incubated in the dark for 1h at room temperature in secondary antibodies (1:500 Goat anti-Guinea Pig, Alexa Fluor® 594 and Goat anti-Mouse Alexa Fluor® 488, Invitrogen), washed 3 times for 5 minutes in PBST, and coversliped with ProLong Gold Antifade Mountant with DAPI (Cat. # P36931, Invitrogen) as the mounting medium. Fluorescent imaging was performed using a Keyence BZ-X710 fluorescence microscope (Keyence America, Itasca, IL) and fluorescent area was quantified with ImageJ software (40).

### 2.11 Statistical Analyses

Statistical analyses were performed in SAS Enterprise Guide 7.1 (SAS Institute Inc., Cary, NC). To assess the effect of genotype within diet group on all dependent variables in our animal studies, we used the mixed model procedure. When statistically significant interactions were found, Tukey’s adjustment for multiple comparisons was used to assess the probability of difference between means. For crossover studies, we conducted paired t-tests to assess differences between saline and glucagon injections within animal. For *ex vivo* lipolysis assays, we conduced paired t-tests to assess differences between control and treatment incubations within each animal. Independent variables were identified as classification variables in all models. Raw data was plotted in Graphpad PRISM® Version 8 for Windows (GraphPad Software, San Diego, California, USA.). All data are presented as mean ± standard error.

## 3. Results

### 3.1 Glucagon signaling does not regulate lipolysis at white adipose tissue

We initially set out to assess the effects of glucagon receptor signaling on adipose tissue lipolysis by examining the effects of acute exogenous glucagon administration on serum NEFA concentration in wildtype C57BL/6J mice. In this time-course study, IP glucagon increased serum glucagon in mice fasted for 4h and 16h compared to saline injected mice at 15 minutes after injection (P<0.05) with a return to levels near baseline by 30 minutes (Figure 1A-B). Glucagon administration had no effect on serum NEFA concentration (Figure 1C-D). Consistent with glucagon’s stimulatory effect on glycogenolysis and gluconeogenesis, IP glucagon induced a robust rise in serum glucose in both 4h and 16h fasted mice (Figure 1E-F), with a return to basal levels by 30 minutes after injection. Acute exogenous glucagon administration did not affect serum insulin (Figure 1G-H).

**Figure 1.**
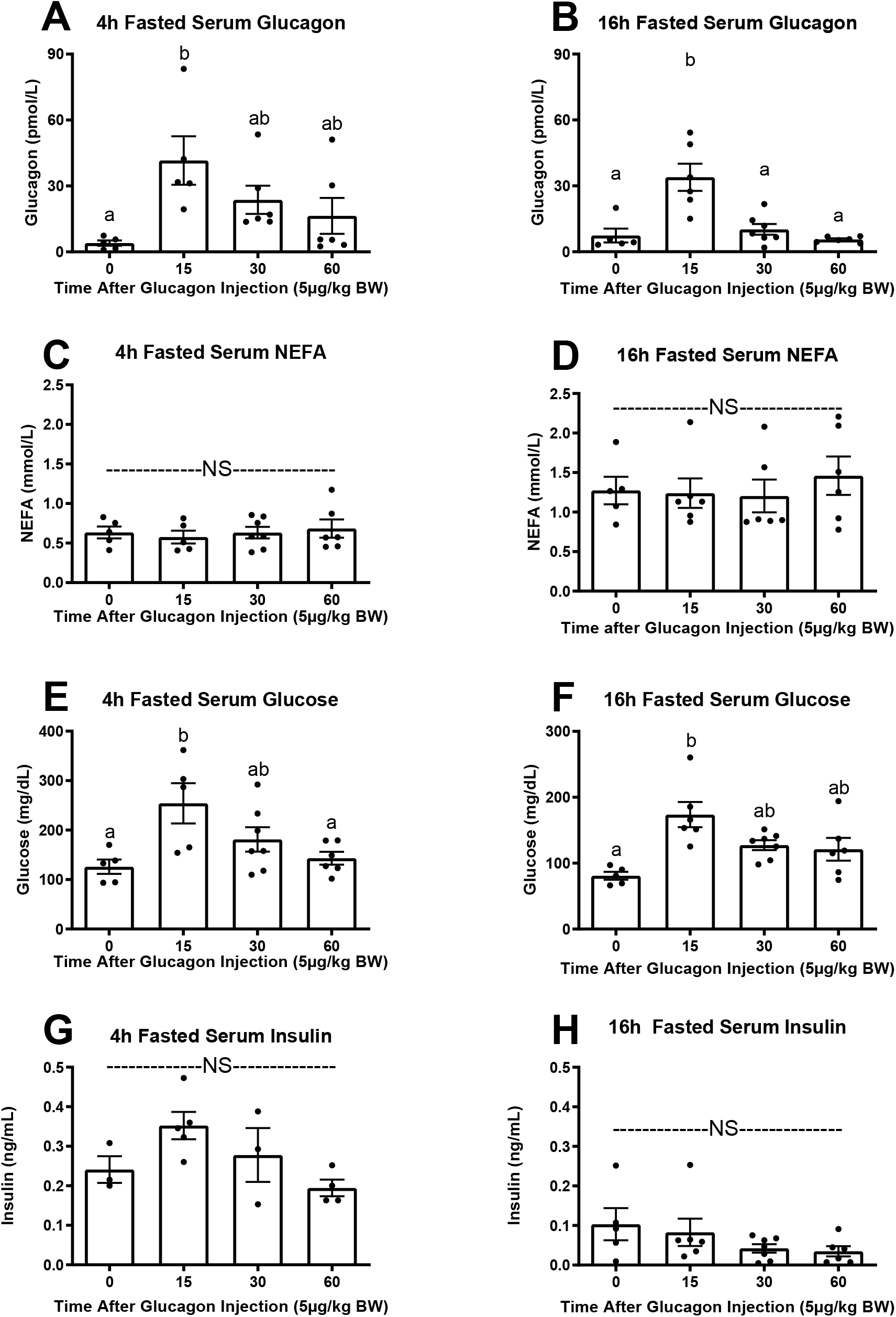
Acute IP glucagon administration in wildtype mice. Serum glucagon, NEFA, glucose, and insulin concentrations in wildtype C57BL/6J mice fasted for 4 (A, C, E, G) or 16 (B, D, F, H) hours (n= 5-7 per group for all except fed insulin, n=3-5 per group). Mice were injected with saline (time 0) or glucagon (5µg/kg) and sacrificed at 15, 30, or 60 minutes after IP injection. Data presented as Mean ± SEM. ^a,b^Superscript letters that differ indicate differences within group, P < 0.05. NS (not significant).

Having demonstrated that exogenous glucagon does not affect circulating NEFA concentration *in vivo*, we set out to examine the direct effects of glucagon on adipose tissue lipolysis in gonadal adipose tissue explants from mice fasted for either 4 or 16h. In explants from 4h fasted wildtype C57BL/6J mice, isoproterenol, which stimulates lipolysis via activation of both β1 and β2 adrenergic receptors, robustly increased NEFA release (P<0.0001, Figure 2A). In rodents, an extended fast increases lipolysis via catecholamine stimulated β adrenergic signaling and decreases the lipolytic response to isoproterenol (41). Accordingly, isoproterenol did not further stimulate lipolysis in explants from mice fasted for 16h (Figure 2B). Forskolin, which stimulates lipolysis by directly activating adenylate cyclase and increasing intracellular cyclic AMP concentration, increased NEFA release in explants collected from 4h and 16h fasted mice (Figure 1A-B; P < 0.01). Insulin (20nM, 200nM, and 200nM) robustly decreases lipolysis in explants from 4h fasted mice (20nM: P=0.012, 200nM: P<0.006, 200nM: P=0.009, Supplemental Figure 7). Consistent with a decrease in insulin’s suppressive action on lipolysis in extended fasting (42), we observed no effect of insulin on NEFA release in explants from 16h fasted mice (Supplemental Figure 7). Varying concentrations of glucagon (0.1nM, 1nM, 10nM, and 100nM) had no effect on NEFA release in explants from mice, regardless of fasting state (Figure 2C-D). We next applied this assay to WAT explants from Gcgr^adipocyte+/+^ and Gcgr^adipocyte-/-^ mice. We found that glucagon receptor expression at WAT did not affect forskolin stimulated lipolysis, regardless of fasting state (Figure 2E-F). Similar to our findings in explants from wildtype mice, glucagon did not affect *ex vivo* lipolysis in explants from Gcgr^adipocyte+/+^ or Gcgr^adipocyte-/-^ mice, regardless of fasting state (Figure 2G-H).

**Figure 2.**
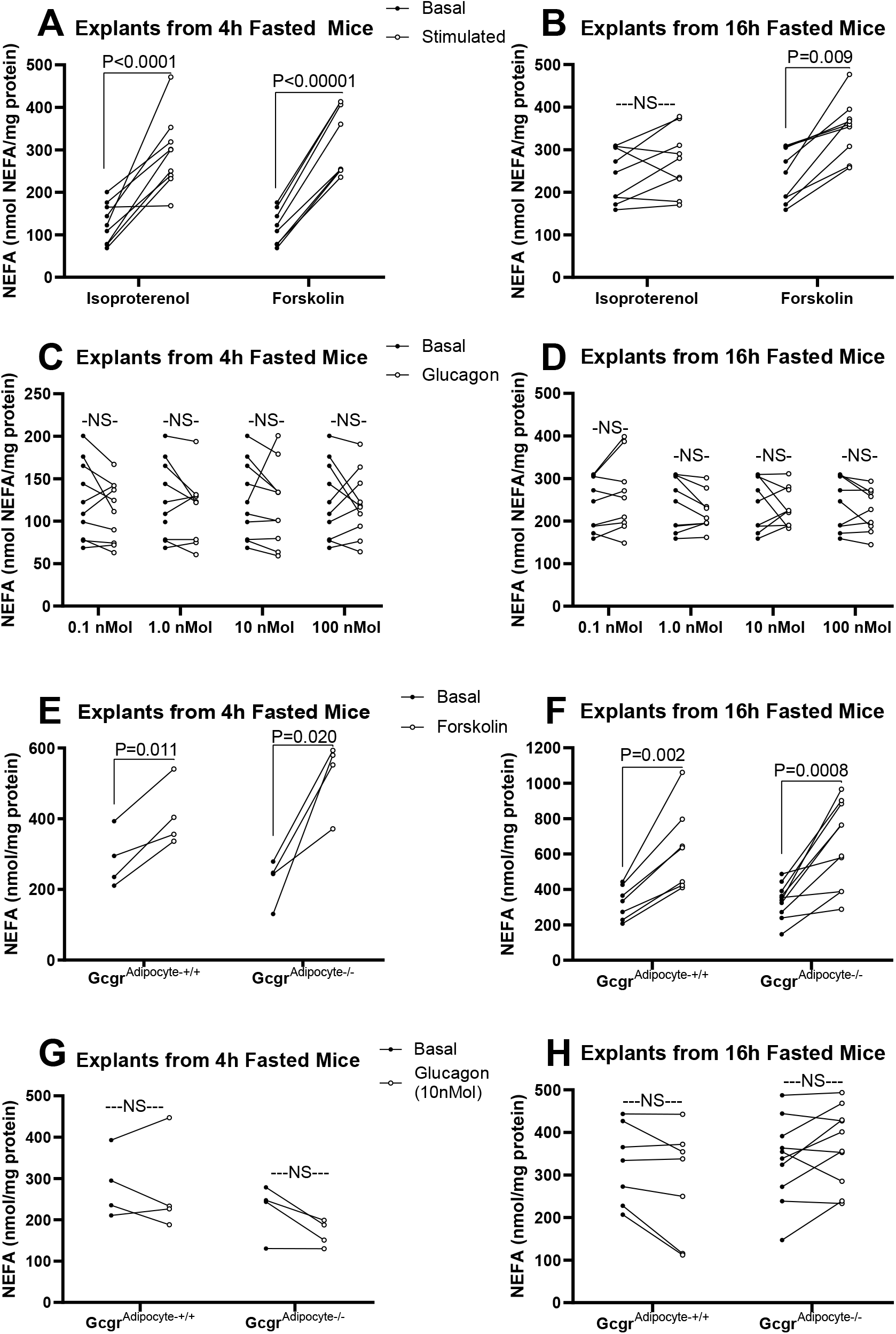
*Ex vivo* lipolysis from gonadal adipose tissue explants. Explant NEFA release in response to bath application of isoproterenol, forskolin and glucagon was assessed in 4h and 16h fasted mice. Isoproterenol and forskolin stimulated explant NEFA release from wildtype C57BL/6J mice fasted for 4h (A; n=10) or 16h (B; n= 8). Media NEFA concentrations induced by glucagon in 4h (C; n=10) and 16h (D; n = 8) fasted wildtype C57BL/6J mice. NEFA release from explants collected from from Gcgr^Adipocyte+/+^ versus Gcgr^Adipocyte-/-^ mice fasted for 4h (n=4) or 16h (Gcgr^adipocyte+/+^: n= 7, Gcgr^adipocyte-/-^: n= 10) and treated with forskolin (E and F) or glucagon (G and H). All studies were performed in triplicate explants from each mouse. Data presented as Mean ± SEM. NS (not significant).

Fasting stimulates lipolysis, increases NEFA release and induces an increase in hepatic triglyceride accumulation (43). Fasting also increases glucagon secretion and signaling (17). To assess if glucagon signaling mediates fasting-induced lipolysis and hepatic lipid accumulation, we examined the effects of an extended fast (16h) on serum NEFA and hepatic triglyceride concentrations in Gcgr^adipocyte+/+^ and Gcgr^adipocyte-/-^ mice. Fasting robustly and equally increased both serum NEFA and hepatic triglyceride concentrations in lean Gcgr^adipocyte+/+^ and Gcgr^adipocyte-/-^ mice (P < 0.01, Figure 3A-B). Similarly, diet induced obese Gcgr^adipocyte+/+^ and Gcgr^adipocyte-/-^ mice respond to a 16h fast with an equally robust rise in serum NEFA concentration and hepatic triglyceride accumulation (P<0.05, Supplemental Figure 3). Acute exogenous IP glucagon did not affect serum NEFA concentration in either Gcgr^adipocyte+/+^ or Gcgr^adipocyte-/-^ mice. However, consistent with the gluconeogenic and glycogenolytic actions of glucagon at the liver, IP glucagon increased serum glucose, regardless of genotype (Gcgr^adipocyte+/+^: P=0.025, Gcgr^adipocyte-/-^: P=0.002, Figure 3C-D).

**Figure 3.**
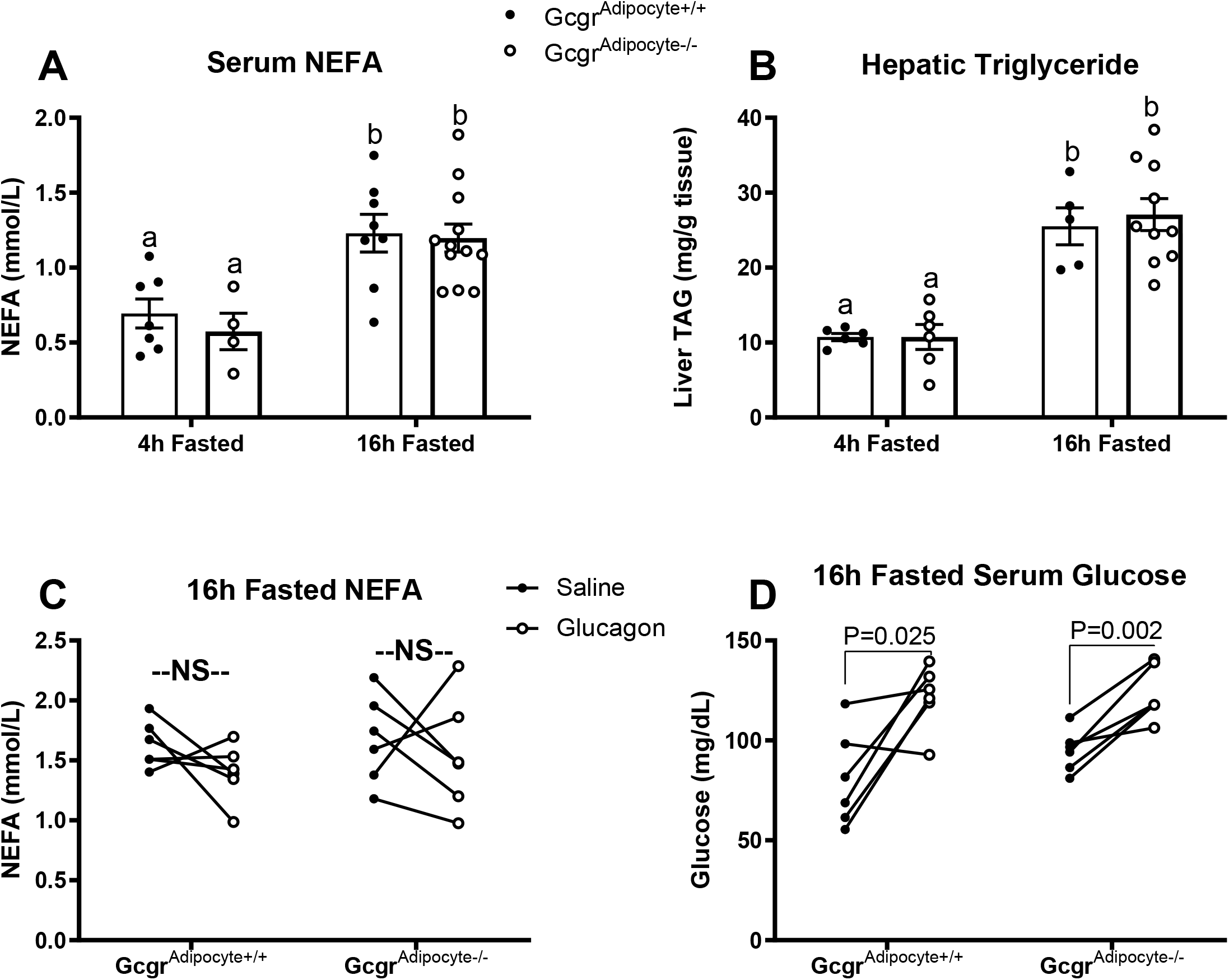
Gcgr^adipocyte-/-^ mice fed a low-fat diet have normal fasting-induced changes in serum NEFA concentration and hepatic lipid accumulation. A) Serum NEFA and B) hepatic triglyceride concentrations in mice fasted for 4h (n= 6-7 Gcgr^adipocyte+/+^ and n=4-6 Gcgr^adipocyte-/-^) and 16h (n= 7 Gcgr^adipocyte+/+^ and n=10-11 Gcgr^adipocyte-/-^). Serum NEFA (C) and glucose (D) concentrations in 16h fasted Gcgr^adipocyte+/+^ and Gcgr^adipocyte-/-^ mice injected with saline and glucagon (n=6 per genotype). ^a,b^Superscript letters that differ indicate differences within group, P < 0.01. Data presented as Mean ± SEM. NS (not significant).

Both fasting (44) and glucagon (3, 45) increase hepatic FGF21 production via PPARα activation. FGF21 stimulates lipolysis in white adipose tissue (44). We found that both Gcgr^adipocyte+/+^ and Gcgr^adipocyte-/-^ mice respond to a 16h fast with an increase in serum FGF21 (P<0.01 for both). Acute IP glucagon administration also increased serum FGF21 in both groups 30 minutes after injection (P=0.03 for Gcgr^adipocyte+/+^ and P=0.01 for Gcgr^adipocyte-/-^, Supplemental Figure 5A-B).

Circulating NEFA concentration is dependent on the balance between fatty acid release by adipose tissue and clearance by other tissues. Lipid clearance after an olive oil oral gavage did not differ between Gcgr^adipocyte+/+^ and Gcgr^adipocyte-/-^ mice (Supplemental Figure 2A-B). Glucagon decreases hepatic triglyceride secretion (2) and chronic glucagon administration decreases serum cholesterol (46, 47). IV injection of the nonionic detergent, Triton WR1339, inhibits triglyceride hydrolysis by lipoprotein lipase, thereby providing an indication of hepatic triglyceride production. Hepatic triglyceride secretion, as assessed by Triton WR1339, did not differ between Gcgr^adipocyte+/+^ and Gcgr^adipocyte-/-^ mice (Supplemental Figure 2C), nor did 4h fasted serum cholesterol in lean (Supplemental Figure 2D) or obese mice (Supplemental Figure 3H).

### 3.2 Glucagon signaling at the adipocyte does not affect adipokine release

Leptin stimulates adipose tissue lipolysis (48), while adiponectin decreases lipolysis (49). Thus, we explored whether glucagon signaling at the adipocyte regulates the release of these adipokine-mediators of lipolysis. Gcgr^adipocyte+/+^ and Gcgr^adipocyte-/-^ mice respond to a 16h fast with an equally robust decrease in serum leptin concentration (P<0.01, Supplemental Figure 4A). Acute IP glucagon administration does not affect serum leptin or adiponectin concentrations (Supplemental Figures 4B and 6A-B) and incubation of adipose tissue explants with glucagon does not affect leptin release into the media (Supplemental Figure 4C).

### 3.3 Glucagon signaling at the adipocyte does not regulate glucose homeostasis

Because glucagon receptor signaling at the liver plays a critical role in the maintenance of glucose homeostasis, we set out to determine if glucagon-receptor signaling at the adipocyte affects glucose homeostasis. Body weight (Figure 4A), oral glucose clearance (Figure 4B-C), glucose-stimulated insulin secretion (Figure 4D), and insulin tolerance (Figure 4E-F) did not differ between Gcgr^adipocyte+/+^ and Gcgr^adipocyte-/-^ mice fed a low-fat diet. Similarly, we observed no differences in diet-induced weight gain, oral glucose clearance, or glucose-stimulated insulin in diet-induced obese Gcgr^adipocyte+/+^ and Gcgr^adipocyte-/-^ mice (Supplemental Figure 3B-D). Reflective of glucagon’s critical role in the liver, mice lacking the glucagon receptor at the hepatocyte are hyperglucagonemic with islets that exhibit severe α-cell hyperplasia (1). In islets from both lean and diet-induced obese mice, α-cell abundance (% of total islet area) did not differ between Gcgr^adipocyte+/+^ and Gcgr^adipocyte-/-^ mice (Figure 5C-H). Furthermore, despite a significant reduction of Gcgr mRNA expression in gonadal WAT, Gcgr^adipocyte-/-^ mice have an equally high expression of hepatic Gcgr mRNA as Gcgr^adipocyte+/+^ mice and similar concentrations of serum glucagon in the fasted state (Figure 5A-B). In line with these findings, 16 hours of fasting equally increased\ (P<0.05) hepatic mRNA expression of Gcgr and the glucagon-responsive genes: Phosphoenolpyruvate Carboxykinase (Pck1), Peroxisome proliferator-activated receptor-alpha (Ppara), and Carnitine Palmitoyl Transferase-1a (CPT1a) in Gcgr^adipocyte+/+^ and Gcgr^adipocyte-/-^ mice, with no differences between genotype within fasting duration (Supplemental Figure 1).

**Figure 4.**
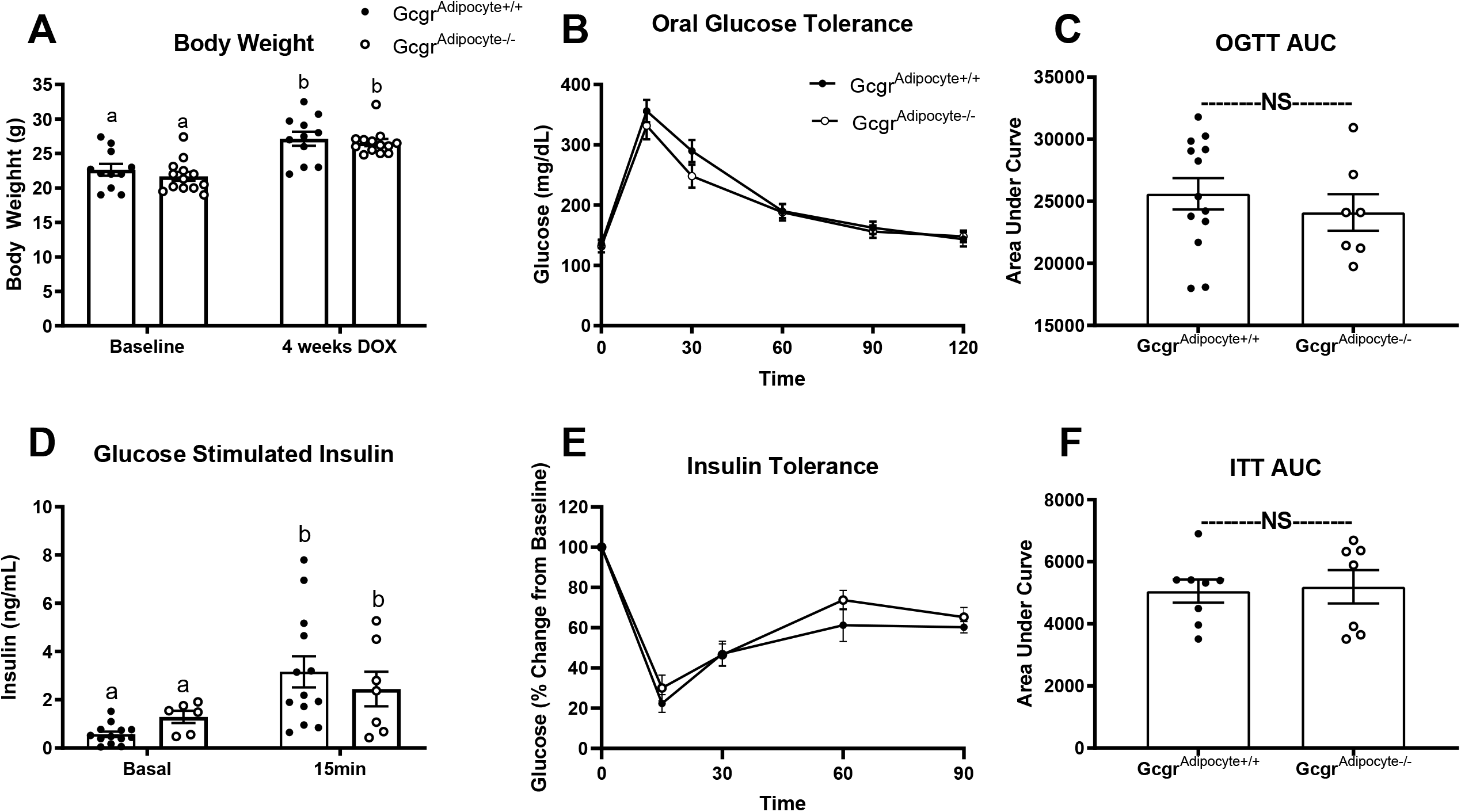
Gcgr^adipocyte-/-^ mice fed a low fat diet have a normal response to oral glucose. A) body weight, B) oral glucose tolerance (OGTT), C) OGTT area under the curve, D) glucose-stimulated insulin, E) insulin tolerance test (ITT), and F) ITT area under the curve in lean mice (For OGTT, glucose-stimulated insulin, and ITT n= 13 Gcgr^adipocyte+/+^ and n=7 Gcgr^adipocyte-/-^ mice. For body weight n= 11 Gcgr^adipocyte+/+^ and n= 13 Gcgr^adipocyte-/-^ mice). ^a,b^Superscript letters that differ indicate differences within group, P < 0.01. NS (not significant). Data presented as Mean ± SEM.

**Figure 5.**
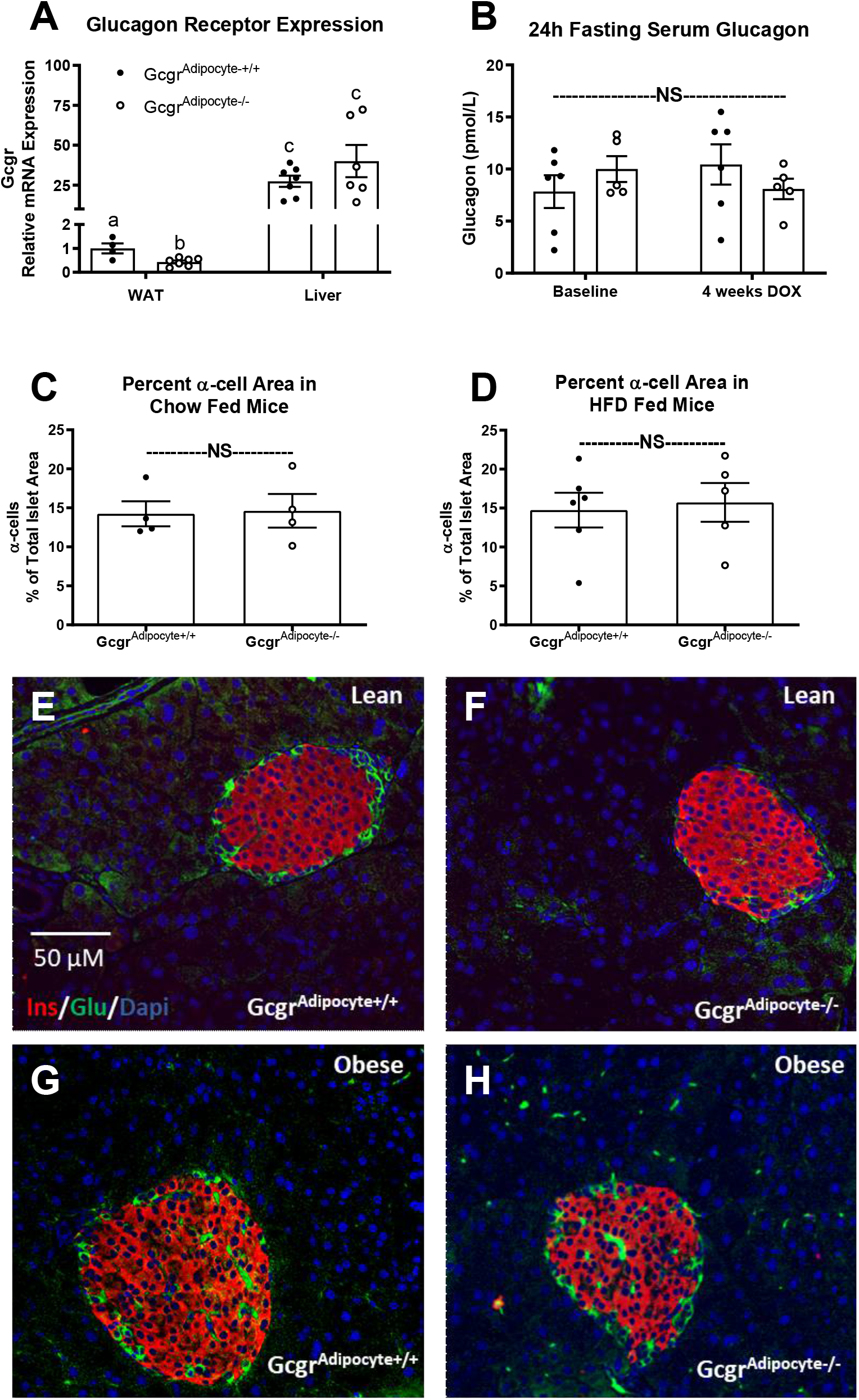
Gcgr gene expression, fasting glucagon, and α-cell abundance in Gcgr^adipocyte+/+^ and Gcgr^adipocyte-/-^ mice. A) Relative mRNA expression of glucagon receptor in liver (n= 7 per genotype) and gonadal adipose tissue (n=4 Gcgr^adipocyte+/+^ and n=7 Gcgr^adipocyte-/-^) in lean mice. B) 24h fasting serum glucagon (B) in lean Gcgr^adipocyte+/+^ (n=6) and Gcgr^adipocyte-/-^ (n=5) mice before and after 4 weeks of doxycycline induction. C) α-cell percentage of total islet area in lean (n=4 per genotype) and D) obese (n= 6 Gcgr^adipocyte+/+^ and n=5 Gcgr^adipocyte-/-^) mice with representative photos (E-H) ^a,b,c^Superscript letters that differ indicate differences within group, P < 0.05. NS (not significant). Data presented as Mean ± SEM.

## 4. Discussion

The potential lipolytic role of glucagon signaling at WAT has long been debated. Glucagon secretion rises in response to an extended fast in lean rodents(17) and humans(50, 51), as does lipolysis(13, 43). Because lipolysis is regulated by G-protein coupled receptors, such as adrenergic receptors which activate protein kinase A and increase intracellular concentrations of cAMP (52), it has often been assumed that glucagon also mediates fasting-induced WAT lipolysis in a similar fashion through its G-protein coupled receptor. Contrary to this notion, we report that glucagon does not affect WAT lipolysis either indirectly *in vivo* or through direct action at white adipose tissue *ex vivo*.

Because fasting stimulates adipose tissue lipolysis through adrenergic signals and increases glucagon secretion, independently, it is difficult to ascertain whether glucagon regulates adipose tissue lipolysis or if the rise in glucagon is simply a concomitant result of fasting. Furthermore, glucagon stimulates intrahepatic lipolysis (53), leading to a rise in circulating NEFA concentration. Our *ex vivo* lipolysis assays eliminate these confounding factors and clarify that glucagon does not exert direct lipolytic effects on WAT.

Few studies have fully examined the effects of glucagon on WAT lipolysis *in vivo* in mice. The majority of studies proposing that glucagon does have a lipolytic effect on WAT were performed on isolated adipocytes from rats (8-12, 54) and humans (6) using supraphysiologic levels of glucagon. The generation of mice with a significant reduction of WAT *Gcgr* expression provided a model with which to further explore the physiological role of glucagon receptor signaling at WAT. Using this model, we found that fasting exerted an equally robust increase of serum NEFA concentration and that exogenous glucagon did not affect serum NEFA concentration, regardless of genotype, confirming that glucagon does not exert physiologically relevant effects on adipose tissue lipolysis *in vivo*. Using this model, we also confirmed that glucagon receptor signaling at WAT does not regulate whole body glucose homeostasis. These studies are in contrast to previous reports that supraphysiological levels of glucagon can increase adipocyte glucose uptake *in vitro* (6). Finally, applying our *ex vivo* lipolysis assays to explants from Gcgr^adipocyte+/+^ and Gcgr^adipocyte-/-^ mice confirmed our finding from experiments in explants from wildtype mice that glucagon does not directly regulate lipolysis in WAT *ex vivo*.

While Arafat et al. showed that IP glucagon administration does increase serum NEFA concentration *in vivo* in streptozotocin-induced insulin deficient mice, suggesting that glucagon may, in fact, stimulate WAT lipolysis, the authors did not include a healthy non-diabetic group of mice with sufficient insulin levels (54). Glucagon stimulates intrahepatic lipolysis (53), which is counteracted by insulin (55). Because these studies were only performed in insulinopenic mice, it is difficult to assess if the increase in serum NEFA concentration was due to unregulated hepatic lipolysis as a result of insufficient insulin-mediated suppression of lipolysis or a true increase in adipose tissue lipolysis. Furthermore, the dosage of IP glucagon administered in the studies of Arafat et al. (0.05 mg/kg body weight) is supraphysiological. If we estimate that blood represents approximately 7% of total body weight (56), this dosage would equate to blood glucagon levels of approximately 200 nmol/L. Serum glucagon ranges from <1-7 nmol/L in a lean fed mouse to ∼10-20nmol/L in a 16-24h fasted lean mouse and can reach as high as ∼30 nmol/L in an obese insulin-resistant mouse in the fed state (17). We employed a dosage of 5µg/kg body weight glucagon which, if diluted in blood, could reach a concentration of approximately 20 nmol/L, a level that is within physiological glucagon levels, yet still high. Clinical studies showing physiological levels of glucagon do not effect on adipose tissue lipolysis in either people with (13) or without (13, 14) diabetes support our findings that glucagon does not regulate WAT lipolysis in mice.

In light of our findings, it is important to note that adipocytes are not the major cell type expressing the glucagon receptor within WAT. In fact, through a series of single cell RNA-seq studies, Campbell and colleagues (57) recently showed that glucagon receptor is predominantly localized to pericytes, with no detectable expression in adipocytes. This was true for both mouse and human WAT. These findings explain why our mouse model of adipocyte targeted Gcgr knockout only decreased Gcgr mRNA by ∼50% in whole WAT.

We recognize that our studies have some limitations. Aberrant glucagon secretion and signaling is a hallmark of both type 2 (insulin resistant) and type I (insulin deficient) diabetes. Our studies focused on the role of WAT glucagon receptor signaling in healthy lean mice and diet-induced obese, insulin resistant mice and our conclusions are limited as such. Glucagon receptor antagonists are in clinical trials as a treatment to lower blood glucose in patients with both type 2 and type1 diabetes. Thus, further exploration into the potential role of WAT glucagon receptor signaling in type 1 diabetes is required to understand the potential impact of the use of glucagon receptor antagonists in this patient population.

## 5. Conclusion

We have established that glucagon does not regulate WAT lipolysis, either directly or indirectly. Further, our studies show that glucagon receptor signaling at WAT does not affect whole body lipid or glucose homeostasis.

## Author Contributions

J.H.S. and J.L.B conceived and designed research. J.H.S., T.J.M., and A.V. performed experiments and data analysis. T.J.M. and A.V. performed *ex vivo* lipolysis assays in wildtype mice. J.H.S. and J.L.B. interpreted results of experiments. J.H.S drafted manuscript. J.L.B., T.J.M., and A.V. edited and revised manuscript.

## Acknowledgements

The authors thank Dr. Daniel Drucker, M.D. for his vital contribution to study design, interpretation of data, and for the use of the floxed Gcgr mouse. The authors thank Philipp Scherer, Ph.D. for providing the Adiponectin-rtTA and TRE-Cre mice. The authors would also like to acknowledge the role of the late Dr. Roger Unger, M.D. for his contribution to study design and interpretation of data. Dr. Unger was a remarkable mentor, colleague, and friend. He is missed dearly.

## Supplementary Figures

**Figure S1.**
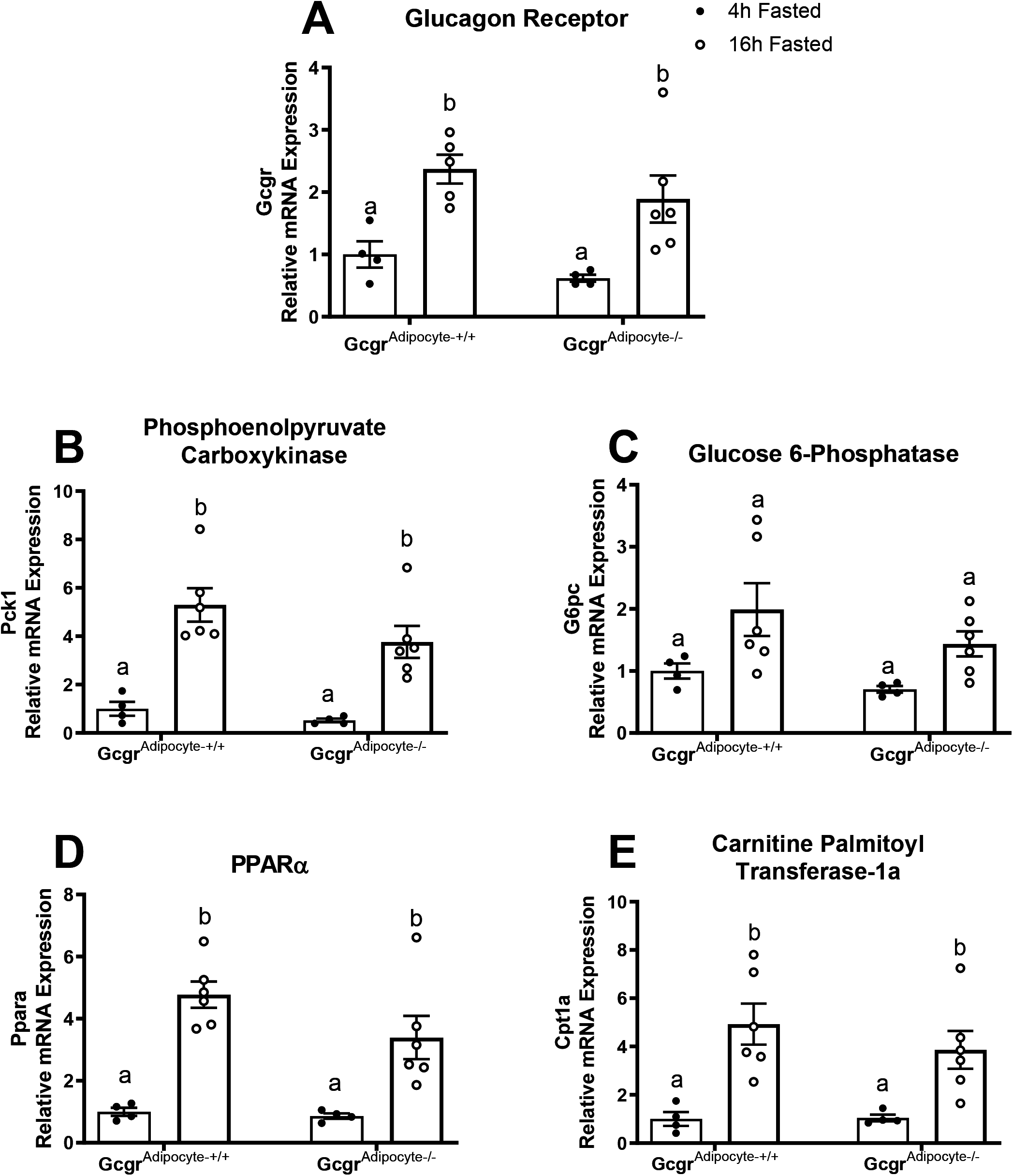
Hepatic expression of mRNA encoding for Gcgr and and glucagon-responsive genes in 4h and 16h fasted Gcgr^adipocyte+/+^ and Gcgr^adipocyte-/-^ mice fed a chow diet. Relative hepatic mRNA expression of A) glucagon receptor (Gcgr) and glucagon-responsive genes: B) Phosphoenolpyruvate Carboxykinase (Pck1), C) Glucose 6-Phosphatase (G6pc), D) Peroxisome proliferator-activated receptor α (Ppara), and E) Carnitine Palmitoyl Transferase-1a (Cpt1a) in 4h (Gcgr^adipocyte+/+^: n=4, Gcgr^adipocyte-/-^: n= 4) and 16h fasted (Gcgr^adipocyte+/+^: n=6, Gcgr^adipocyte-/-^: n= 6) mice. Gene expression was normalized to the housekeeping gene ACTβ and presented as relative expression compared to the Gcgr^adipocyte+/+^ 4h fasted group. ^a,b^Letters that differ indicate differences within group, P < 0.05. Data presented as Mean ± SEM.

**Figure S2.**
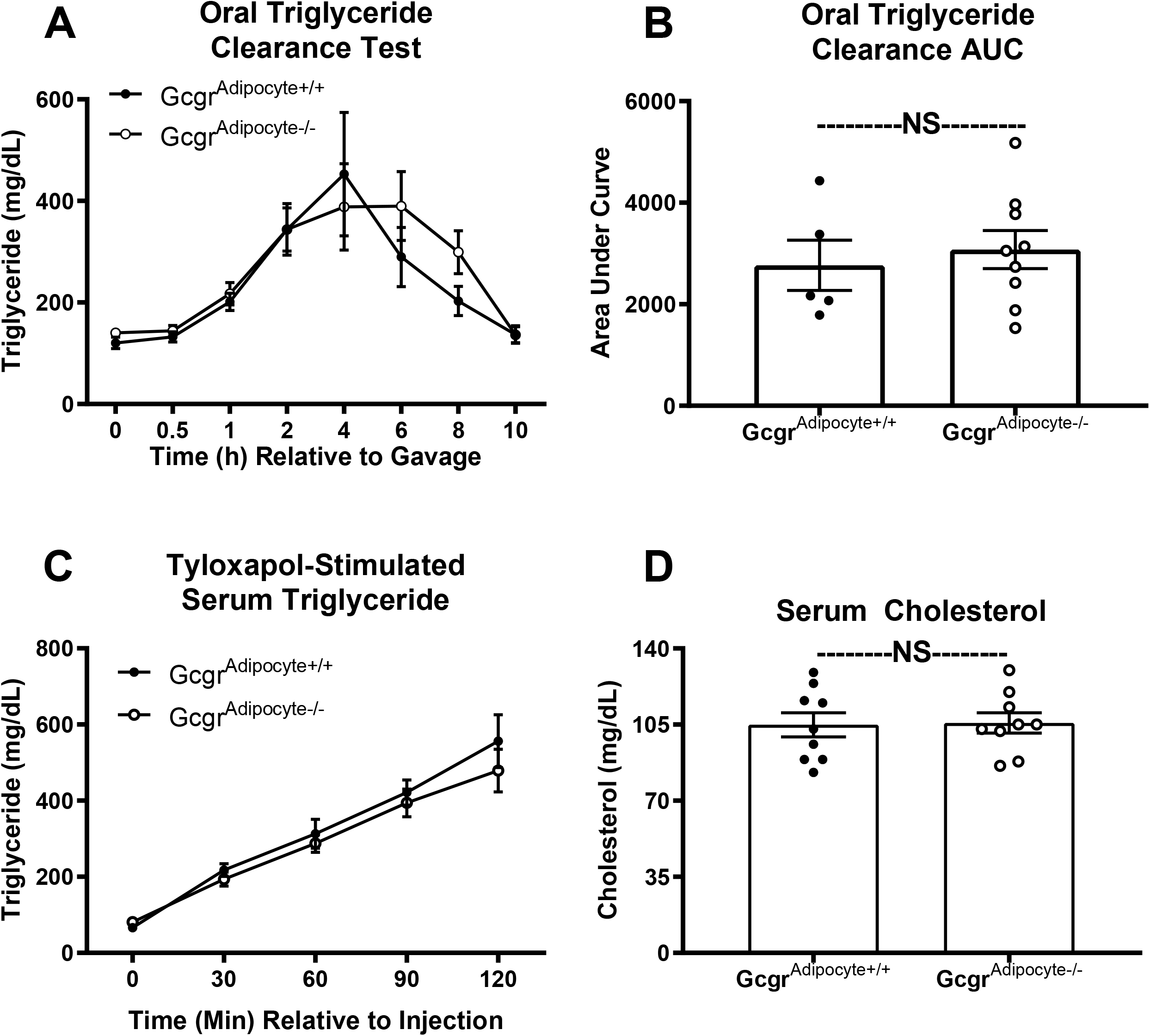
Lean Gcgr^adipocyte-/-^ mice have normal lipid clearance, hepatic triglyceride secretion, and serum total cholesterol. Oral lipid clearance in mice gavaged with olive oil (10 µL/g body weight) expressed as a A) time course and B) area under the curve (AUC); Gcgr^adipocyte+/+^: n=6, Gcgr^adipocyte-/-^: n= 9). C) Tyloxapol stimulated serum triglyceride concentration (n=6 per genotype). D) 4h fasted serum cholesterol in lean mice (n=9 per genotype). NS (not significant). Data presented as Mean ± SEM.

**Figure S3.**
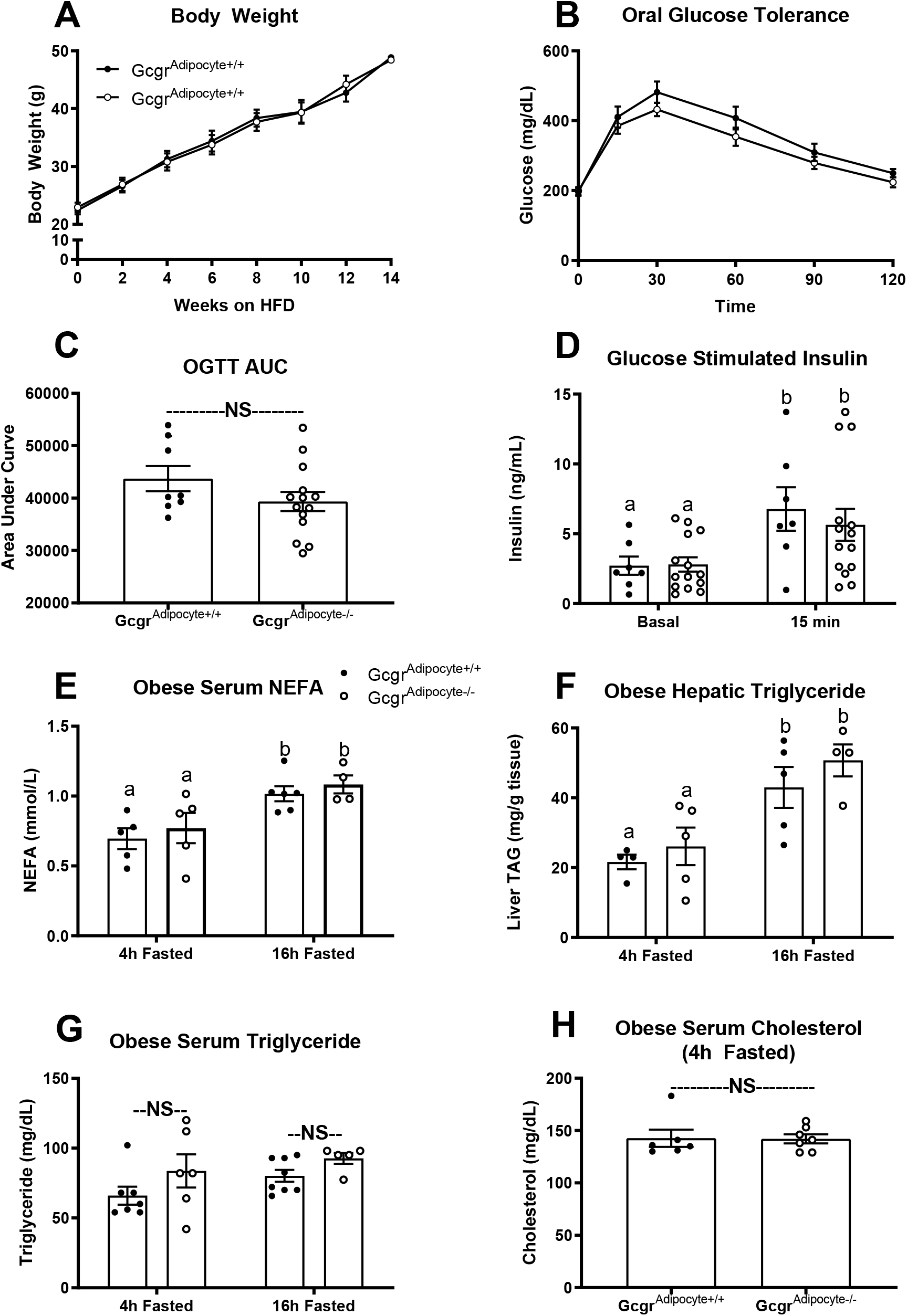
Diet-induced obesity similarly affects metabolism in Gcgr^adipocyte+/+^ and Gcgr^adipocyte-/-^ mice. A) Body weight gain in mice provided a high fat diet for 14 weeks (Gcgr^adipocyte+/+^: n=12, Gcgr^adipocyte-/-^: n= 13). B) Oral glucose tolerance, C) OGTT area under the curve, and D) oral glucose-stimulated serum insulin concentration (Gcgr^adipocyte+/+^: n= 7-8, Gcgr^adipocyte-/-^: n= 14) after 14 weeks on a high-fat diet. Serum E) NEFA concentration (Gcgr^adipocyte+/+^: n= 6, Gcgr^adipocyte-/-^: n= 5), F) hepatic triglyceride concentration (Gcgr^adipocyte+/+^: n= 4 for 4h fasted and n=5 for 16h fasted, Gcgr^adipocyte-/-^: n= 5 for 4h fasted and n=4 for 16h fasted), G) serum triglyceride concentration (Gcgr^adipocyte+/+^: n= 7 for 4h fasted and n=8 for 16h fasted, Gcgr^adipocyte-/-^: n= 6 for 4h fasted and n=5 for 16h fasted), and H) serum cholesterol concentration (Gcgr^adipocyte+/+^: n= 6, Gcgr^adipocyte-/-^: n= 7) in diet-induced obese mice. ^a,b^Superscript letters that differ indicate differences within group, P < 0.01. NS (not significant). Data presented as Mean ± SEM.

**Figure S4.**
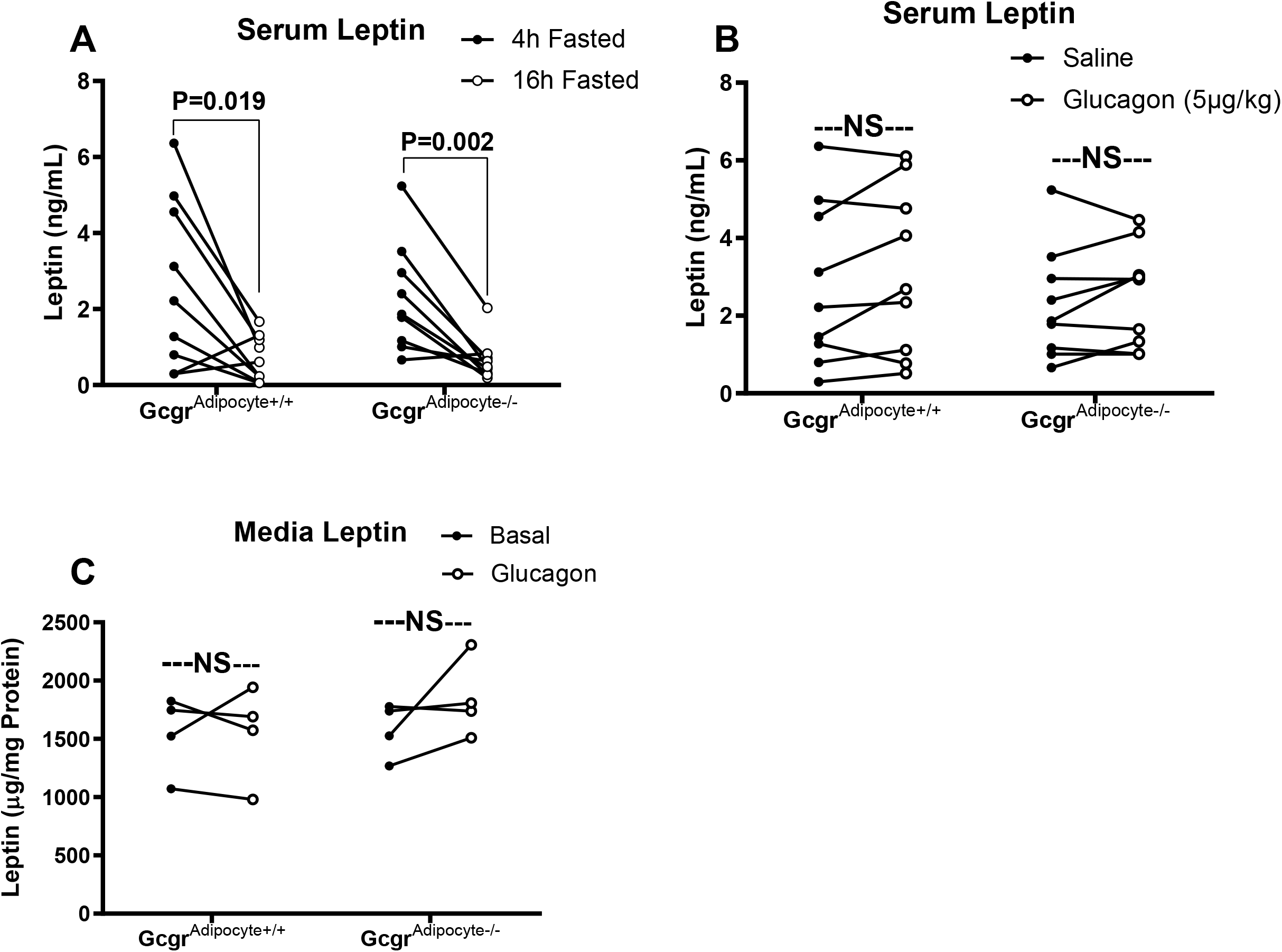
Glucagon receptor signaling does not regulate adipose tissue leptin secretion. A) Serum leptin concentration in mice fasted for 4h and 16h (n=9 per genotype), B) serum leptin in 4h fasted mice injected with saline or glucagon (5µg/kg) (n=9 per genotype), and C) media leptin from gonadal adipose tissue explants (n= 4 mice per genotype in triplicate) incubated in either a control media (Krebs-Ringer Buffer) or media containing glucagon (10nM glucagon) for 1 hour. NS (not significant).

**Figure S5.**
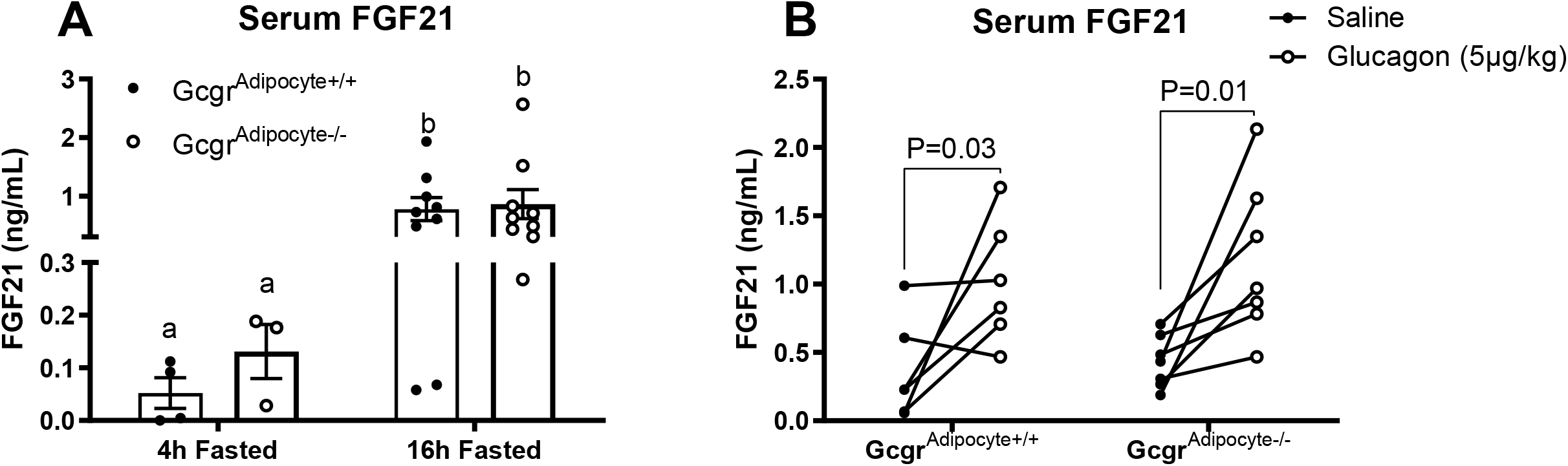
Fasting and glucagon-stimulated FGF21 is normal in Gcgr^adipocyte-/-^ mice fed a low-fat diet. Serum FGF21 in A) mice fasted for 4h (Gcgr^adipocyte+/+^: n= 4, Gcgr^adipocyte-/-^: n= 3), and 16h (n= 9 per genotype) and B) 16h fasted mice injected with saline or glucagon (5µg/kg) (Gcgr^adipocyte+/+^: n= 6, Gcgr^adipocyte-/-^: n= 7). ^a,b^Superscript letters that differ indicate differences within group, P < 0.01. Data presented as Mean ± SEM.

**Figure S6.**
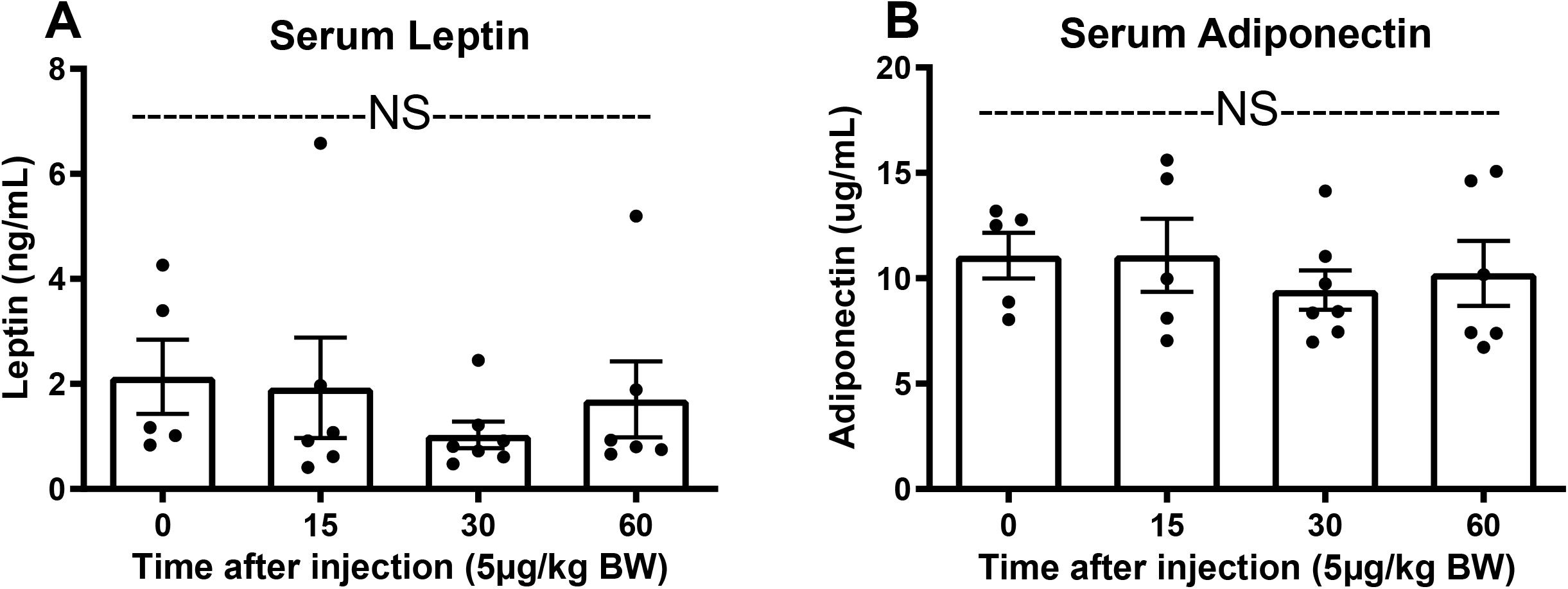
Acute exogenous glucagon administration does not affect serum leptin or adiponectin. Serum A) leptin and B) adiponectin in wildtype mice fasted for 4h (n= 5-7 per group) and injected with saline (time 0) or glucagon (5µg/kg) and sacrificed 0, 15, 30, or 60 minutes after IP injection). NS (not significant). Data presented as Mean ± SEM.

**Figure S7.**
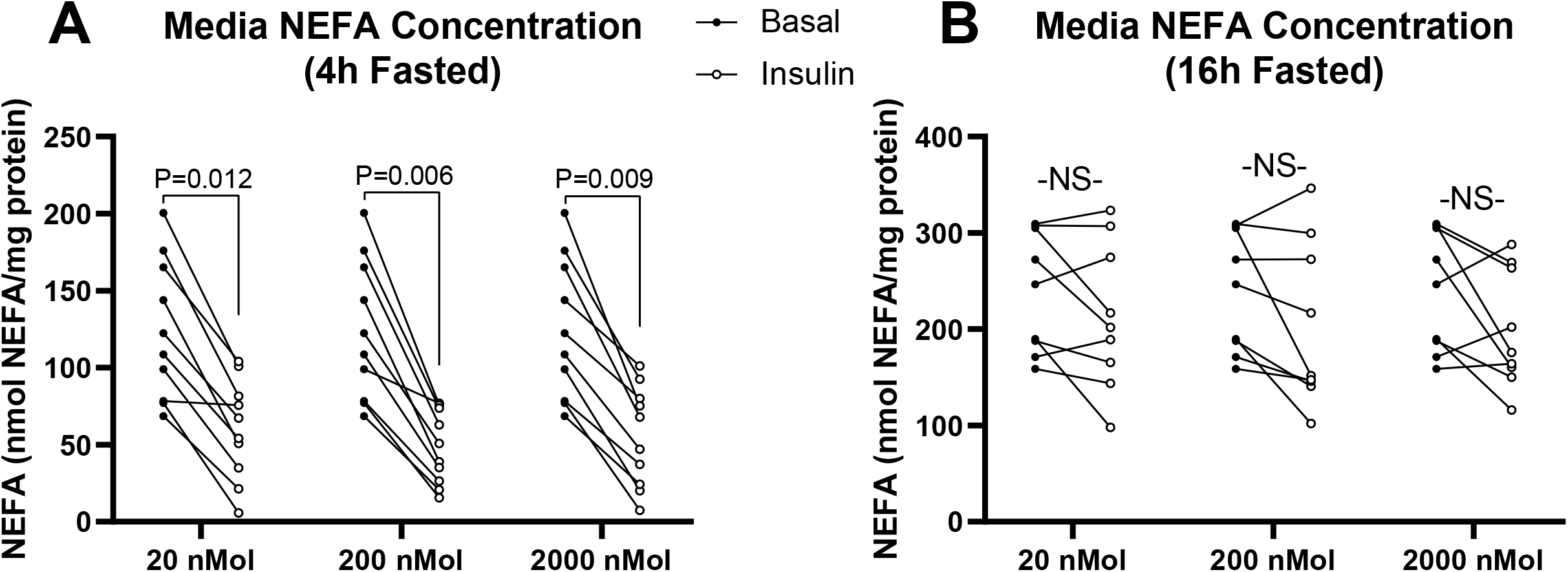
Insulin inhibits lipolysis in gonadal adipose tissue explants from mice fasted for 4h but not 16h. *Ex vivo* lipolysis in explants from wildtype mice that we fasted for A) 4h (n=10) and B) 16h (n=8). Triplicate explants from each mouse were incubated in control media (Krebs-Ringer Buffer; filled circles) and 20nM, 200 nM, and 2000 nM (open circles) insulin diluted in Krebs-Ringer Buffer. NS (not significant).

